# Resting-state EEG alpha-BOLD coupling spatially follows cortical cell-type and receptor gradients

**DOI:** 10.64898/2026.04.14.718407

**Authors:** Stanislav Jiricek, Vincent S.C. Chien, Helmut Schmidt, Vlastimil Koudelka, Radek Marecek, Dante Mantini, Jaroslav Hlinka

## Abstract

The coupling between electroencephalography (EEG) and blood-oxygen-level-dependent (BOLD) signals has been investigated across numerous studies, but its neurobiological underpinnings remain poorly understood. Resting-state EEG alpha-BOLD coupling follows a characteristic spatial pattern, shifting from negative correlations in sensory regions to positive correlations in association cortices. In this study, we examined neurobiological correlates of resting-state alpha-BOLD coupling. We compared the spatial pattern of the alpha-BOLD coupling map to 82 cortical feature maps, including gene expression profiles of different cell types and receptor subunits as well as structural MRI measures. We identified three statistically significant (*q <* 0.05 FDR-corrected) maps: the layer 6 VIP interneuron marker, excitatory layer-5 marker, and NMDA receptor subunit GRIN2C. The three significant gene maps, combined in a multiple linear regression model, explained *R*^2^ = 0.312 of the spatial variance in alpha-BOLD coupling. Analysis of the spatial mismatch between cortical maps and the alpha-BOLD coupling map revealed that the early auditory cortex is the region that consistently diverges from predictions across gene expression and T1/T2 maps. The spatial correspondence between alpha-BOLD coupling and gene expression profiles of specific receptor subunits, neuronal types, and layer-specific populations identifies these as concrete candidates for future computational and experimental studies of alpha-BOLD coupling.

**Author Summary:** The brain’s electrical rhythms and metabolic activity are coupled, yet why this coupling differs across brain regions remains poorly understood. This study shows that resting-state alpha-BOLD coupling, a well-established link between EEG alpha oscillations and fMRI signals, maps onto the brain’s cellular landscape: regions enriched in specific inhibitory interneurons and NMDA receptor subunits show systematically different coupling strengths. These findings suggest that regional differences in cell-type composition and receptor expression, rather than purely anatomical features, could shape the spatial organization of alpha-BOLD coupling. By identifying candidate cortical features, this work can guide future experimental and computational studies, ultimately helping to establish alpha-BOLD coupling as a relevant biomarker for psychiatric and neurological disorders.

## Introduction

For over two decades, simultaneous multimodal neuroimaging has enabled direct study of the relationship between electroencephalography (EEG) and the functional magnetic resonance imaging (fMRI) blood-oxygen-level-dependent (BOLD) signal, yet these relationships remain poorly understood. Few robust EEG-fMRI correlates exist (Moosmann et al., 2003; Laufs et al., 2003a; Mantini et al., 2007; Scheeringa et al., 2008), and clinical applications remain limited (Michels et al., 2021). Here we focus on EEG alpha-BOLD coupling, the most studied resting-state EEG-BOLD correlate (Moosmann et al., 2003; Laufs et al., 2003a; Jiricek et al., 2025). Convergent evidence suggests that, irrespective of how alpha oscillation envelopes are derived, their linear relationship to BOLD follows a unimodal-to-transmodal cortical gradient (Margulies et al., 2016), shifting from negative correlations in predominantly occipital cortex to positive correlations in the default mode network and other associative cortices (Jiricek et al., 2025; Moosmann et al., 2003). Despite this robust pattern, the mechanistic basis of alpha-BOLD coupling remains unclear, limiting its utility for cognitive and clinical research. Understanding this mechanism would enable more principled interpretation of EEG-fMRI data and support the development of biologically grounded computational models.

We hypothesize that this robust spatial heterogeneity of resting-state alpha-BOLD correlation reflects the structural organization of the cortex, specifically that cortical heterogeneities, such as cell-type composition, laminar profiles, and receptor densities, are associated with these correlations.

Cortical gene-expression maps (Gryglewski et al., 2018), primarily derived from the Allen Human Brain Atlas (Hawrylycz et al., 2012), now enable direct comparison with neuroimaging data using established tools and protocols (Markello et al., 2022; Giacomel et al., 2022; Arnatkeviciute et al., 2019; Selvaggi et al., 2021). These spatial maps reveal specific cortical properties, including gene expression profiles and cytoarchitecture. Previous studies have compared spatial features between EEG and fMRI, such as functional connectivity matrices (Wirsich et al., 2021) or structure-function relationships via structural connectivity (Liu et al., 2023), providing insights into brain organization. Relating neuroimaging features to gene expression maps or MRI structural measures has also proven successful in MEG (Mahjoory et al., 2020) and fMRI (Liu et al., 2022) studies, where it allowed formulation of mechanistic hypotheses about brain function. However, whether the spatial pattern of alpha-BOLD coupling itself reflects underlying cortical organization, at the level of cell types, receptor subunits, or laminar profiles, remains poorly understood.

Here, we investigate spatial relationships between the alpha-BOLD coupling map and 82 cortical feature maps (Burt et al., 2018), spanning receptor subunit densities, cell-type gene expression markers, laminar profiles, and MRI-derived structural measures, to identify molecular and anatomical correlates of its regional variability. This analysis is not mechanistic in itself, but generates biologically grounded candidates to inform and constrain future computational models aimed at testing explicit mechanistic hypotheses.

## Methods

### Experimental design and EEG-fMRI data acquisition

We used the group-average statistical map from our previous study (Jiricek et al., 2025) to represent EEG alpha-BOLD coupling. Full methodological details are provided therein. We analyzed simultaneous EEG-fMRI recordings from 72 healthy participants (mean age: 31.4 years, range: 18.3–50.6; 36 males, 36 females), constituting one of the largest simultaneous EEG-fMRI datasets reported to date. The study was approved by the ethics committee of Masaryk University, conducted in accordance with the Declaration of Helsinki, and all participants provided written informed consent. Data were collected at the Central European Institute of Technology (CEITEC) in Brno, Czech Republic. Each participant underwent a 20-minute resting-state session with eyes closed while EEG and fMRI signals were simultaneously recorded. Functional MRI data were acquired on a 3 T Siemens Prisma scanner (64-channel RF head coil) using a multiband, multi-echo EPI sequence (multiband factor = 6; TR = 650 ms; TE = 14.60/33.56/52.52 ms; FA = 30°; 48 axial slices; slice thickness = 3 mm; matrix = 64 *×* 64; FOV = 194 *×* 194 mm; 1,840 volumes). EEG was recorded using a 256-channel MR-compatible EGI HydroCel Geodesic Sensor Net (EGI GES 400 amplifier), sampled at 1000 Hz, along with physiological signals including ECG and respiration.

### EEG-fMRI data preprocessing

Functional MRI preprocessing followed established protocols using SPM12 (Penny et al., 2011) and custom scripts. Steps included spatial realignment, echo fusion via temporal signal-to-noise ratio (tSNR)-weighted averaging, artifact removal using RETROICOR (Glover et al., 2000) for physiological signals, and spatial smoothing with a Gaussian kernel (FWHM = 5 mm). Additional corrections involved regressing potential confounds (24 head motion parameters, first PCA components of white matter and CSF signals) from the gray matter BOLD signal and applying a low-pass filter (cut-off = 0.09 Hz).

EEG preprocessing was conducted using an automated pipeline (Marino et al., 2019; Liu et al., 2017, 2018) in MATLAB, incorporating toolboxes such as SPM (Penny et al., 2011), FieldTrip (Oostenveld et al., 2011), and EEGLAB (Delorme and Makeig, 2004). Gradient artifacts were removed with FASTR (Niazy et al., 2005), while ballistocardiogram artifacts were corrected using the adaptive optimal basis set approach (Marino et al., 2018). Channels with low correlation or high variance in non-physiological frequency bands were identified as noisy and interpolated using the signal of neighboring electrodes.

Further EEG preprocessing involved bandpass filtering (1–80 Hz) and artifact removal through ICA (FastICA). Components corresponding to eye movement (EOG) and muscle activity (EMG) were identified using metrics such as correlation with reference signals, power spectrum characteristics, and kurtosis. Data were subsequently re-referenced to the average reference.

#### Generation of the alpha-BOLD coupling map

We generated the alpha-BOLD coupling map using methodology from our previous study (Jiricek et al., 2025). Briefly, we performed a group-level spatio-spectral decomposition of source-localized EEG data. EEG source localization used individual 12-compartment finite-element head models (SimBio solver) with eLORETA inverse algorithm on a 6 mm source grid. This approach involved the extraction of band-limited power (BLP) for five canonical frequency bands (Delta, Theta, Alpha, Beta, Gamma) and each source position. Independent Component Analysis (ICA) was then applied to all concatenated (along source positions and frequency bands) BLP time series to identify spatio-spectral independent components. Resulting BLP time series was convolved with the hemodynamic response function (HRF) and its derivatives to serve as a regressor in a General Linear Model (GLM). This yielded a group-averaged statistical map (Figure 1, top left) representing the alpha EEG-fMRI pattern and reflecting the voxel-wise relationship between parieto-occipital alpha oscillations and BOLD activity. We leveraged the unthresholded version of this map to transform it to the same space as the maps of the cortical maps data set (Burt et al., 2018). For that purpose, we overlaid our statistical map with the Maximum Probability Map (MPM 1.0) symmetrical brain map (1 mm isotropic resolution) https://identifiers.org/neurovault. image:30759, which indexes the 180 brain areas of the HCP MMP 1.0 atlas (Glasser et al., 2016) in Montreal Neurological Institute (MNI) standard space. We interpolated the values from the unthresholded statistical map (regular grid in gray matter of size 6 mm) to this map by the nearest neighbor interpolation method, and subsequently, all the values belonging to a given brain area were averaged. Note that the HCP MMP 1.0 atlas defines 180 cortical areas per hemisphere (360 total); corresponding areas from the left and right hemispheres were merged to yield 180 bilateral parcels, ensuring compatibility with the single-hemisphere resolution of the human brain transcriptome atlas Hawrylycz et al. (2012) used in Burt et al. (2018) study. This reduction was further supported by the inter-hemispheric symmetry of the EEG alpha-BOLD map (Figure 1, top left).

**Figure 1.**
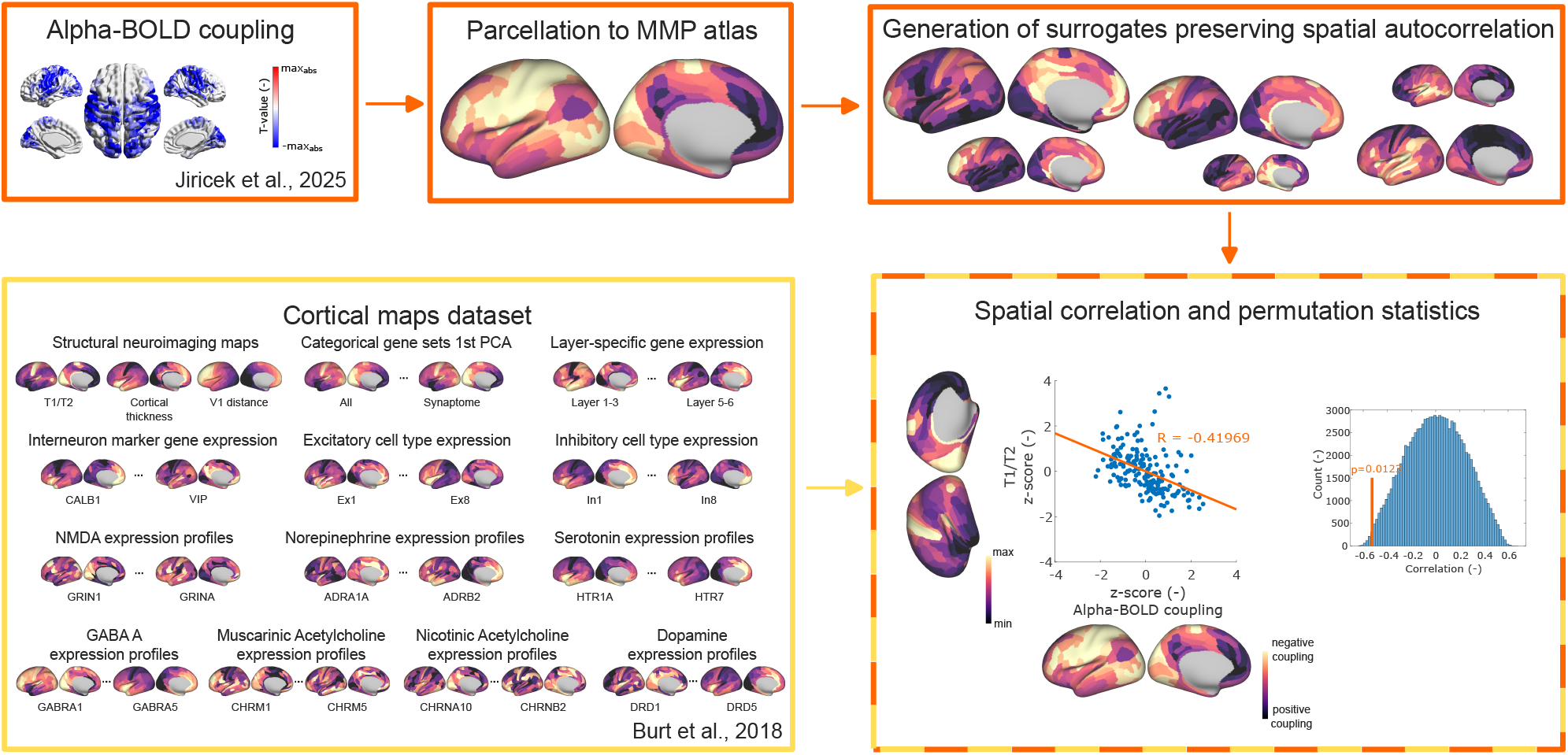
Schematic flowchart of the spatial comparison approach. **Top:** The group-level EEG alpha-BOLD coupling statistical map (voxel-wise *t*-map) is parcellated to the 180-region HCP MMP 1.0 atlas by nearest-neighbor interpolation and parcel-wise averaging. Spatial surrogates of this vector are generated using the BrainSMASH variogram-matching approach, preserving the empirical spatial autocorrelation structure of the original map. **Bottom left:** The cortical maps dataset from Burt et al. (2018) provides 82 parcellated cortical feature maps spanning gene expression profiles of inhibitory and excitatory cell types, receptor subunit densities, laminar-specific expression, and MRI-derived structural measures (T1/T2 ratio, cortical thickness, V1 geodesic distance), all defined on the same HCP MMP 1.0 parcellation. **Right:** For each of the 82 cortical maps, spatial similarity with the alpha-BOLD coupling map is quantified by Spearman correlation. Statistical significance is evaluated against the surrogate null distribution of correlations obtained from the surrogates, and the resulting 82 p-values are subjected to Bonferroni and FDR multiple-comparison corrections.

### Cortical maps dataset

We used a publicly available dataset (https://balsa.wustl.edu/study/Kx5n) of 82 cortical maps from Burt et al. (2018), all parcellated to the same 180-region HCP MMP 1.0 atlas (Glasser et al., 2016), enabling direct spatial comparison with the alpha-BOLD coupling map. Despite sharing the same parcellation resolution, the maps span a wide range of biological scales, from molecular receptor subunit densities, through cellular gene expression markers of specific inhibitory and excitatory neuronal types and laminar populations, to macroscale MRI-derived structural measures, making this dataset well-suited for exploratory, hypothesis-free analysis (Figure 1, bottom left).

Specifically, at the molecular scale, the dataset includes expression profiles of receptor subunits for major neurotransmitter systems: NMDA, GABA, muscarinic and nicotinic acetylcholine, norepinephrine, dopamine, and serotonin. At the cellular scale, it provides gene expression maps distinguishing inhibitory interneuron subtypes and weighted profiles of excitatory and inhibitory neuronal cell types derived from single-cell RNA sequencing. At the mesoscale, laminar-specific expression maps (supragranular, granular, infragranular) and aggregated first principal component (PC1) profiles for predefined gene sets are included. Finally, at the macroscale, three MRI-derived structural maps are provided: T1/T2 contrast, cortical thickness, and V1 geodesic distance.

### Statistical analysis

All statistical comparisons operate on the 180 bilateral HCP MMP 1.0 parcels as the unit of analysis, not on individual participants. Following the methodology of Burt et al. (2018), we evaluated spatial similarity between the alpha-BOLD coupling map and each of the cortical maps using Spearman’s correlation, yielding 82 correlation values. Both map types exhibit strong spatial auto-correlation, complicating significance testing: classical approaches (parametric or standard permutation tests) that ignore spatial autocorrelation inflate false-positive rates (Markello and Misic, 2021). However, several methods were recently designed to tackle this challenge by generating spatially constrained surrogate null distributions (Váša et al., 2018; Vázquez-Rodríguez et al., 2019; Baum et al., 2020; Burt et al., 2020). We used the parametric generative approach from Burt et al. (2020), which creates surrogates matching the spatial autocorrelation structure of a given map via variogram-based modeling (Viladomat et al., 2014). We generated 100,000 surrogate maps of the EEG alpha-BOLD coupling map using the BrainSMASH tool (Burt et al., 2020). The parameter *nh*, which controls the number of uniformly spaced distance bins used to compute the variogram, was increased from the default value of 25 to 50 to ensure that the original map’s variogram fell within the distribution of surrogate variograms. For each cortical gene-expression map, we generated a null distribution of its correlation with the surrogates of alpha-BOLD coupling map. The p-value for each gene map was then determined as the percentile of its empirical correlation within the corresponding surrogate null distribution (two-sided test). The resulting 82 p-values were then subjected to three separate multiple-comparison corrections: Bonferroni (*α*_*BF*_ = 0.05), false discovery rate (*α*_*FDR*_ = 0.05) (Benjamini and Hochberg, 1995), and, in addition, an uncorrected significance threshold (*α*_*uncorr*_ = 0.01) was reported.

#### Explainability analysis

To assess how well the statistically significant gene maps jointly account for the spatial pattern of alpha-BOLD coupling, we constructed a multiple linear regression model (gene-based regression model). The dependent variable (alpha-BOLD coupling) and the three predictor maps that survived FDR correction (*In3, GRIN2C*, and *Ex5*) were each transformed to ranks prior to ordinary least-squares regression. Model fit was quantified by the coefficient of determination *R*^2^. Statistical significance was assessed using the same 100,000 BrainSMASH surrogate maps. For each surrogate, the rank-transformed surrogate map was regressed onto the same rank-transformed predictors and *R*^2^ was computed, yielding a surrogate null distribution. A one-sided p-value was computed as the proportion of surrogates with *R*^2^ exceeding the observed value. Regression residuals (observed minus fitted rank) follow the same sign convention: positive residuals (red) indicate regions where alpha-BOLD coupling is more positive than the model predicts, and negative residuals (blue) indicate regions where it is more negative than predicted. We also constructed an analogous model based on three MRI-derived structural maps as predictors: T1/T2 ratio (a proxy for cortical myelination), cortical thickness, and V1 geodesic distance (the only three structural maps in the Burt et al. (2018) dataset), collectively referred to as the structure-based regression model. The identical rank-transformation, ordinary least-squares fitting, and surrogate-based significance testing procedure was applied, enabling direct comparison of *R*^2^ and p-values between the gene and structural models.

#### Spatial mismatch analysis

To identify which brain regions deviate most from the overall spatial relationship, we computed signed rank differences at the level of individual HCP MMP 1.0 parcels. For each of the 82 cortical maps, both the alpha-BOLD coupling vector and the cortical map vector were converted to ranks via rank transformation. Ranks range from 1 (lowest value) to 180 (highest value), corresponding to the 180 HCP MMP parcels. When the observed Spearman correlation was negative, the cortical map rank vector was reversed to ensure directional consistency across maps. The signed rank difference for parcel *i* was then

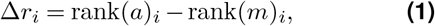

where *a* is the alpha-BOLD coupling vector and *m* the cortical map vector. Positive Δ*r*_*i*_ values (shown in red on brain surface maps) indicate parcels where alpha-BOLD coupling is more positive than predicted by the cortical map. Negative values (blue) indicate parcels where alpha-BOLD coupling is more negative than predicted. Large absolute values |Δ*r*_*i*_| indicate parcels where the two maps disagree most in relative rank position regardless of direction. To assign each parcel to a functional network, the seven canonical resting-state network atlas of Yeo et al. (2011) was resampled onto the MMP 1.0 voxel grid using nearest-neighbor interpolation, and each parcel was assigned to the network occupying the largest number of voxels within its mask. Mean absolute rank difference was then computed within broad anatomical cortex regions defined in the HCP MMP 1.0 atlas (Glasser et al., 2016) and within these functional network assignments, and visualized as bar charts sorted by descending mean rank difference. These rank difference maps are exploratory and descriptive; no inferential testing was applied at the individual-parcel level.

### Data and code accessibility

The raw and preprocessed EEG-fMRI data will be made available by the authors upon reasonable request. The analysis scripts and precomputed features supporting the results reported in this study are publicly available at https://github.com/cobragroup/AlphaMap.

## Results

We compared the alpha-BOLD coupling map to 82 cortical maps from Burt et al. (2018), spanning the smallest to larger cortical scales: receptor subunits, inhibitory and excitatory cell markers, laminar profiles, and MRI-derived structural measures.

### Three cortical maps significantly correlate with alpha-BOLD coupling after FDR correction

We identified three statistically significant spatial correlations of the alpha-BOLD coupling map with interneuron marker VIP (In3 cluster, Bonferroni corrected), excitatory layer-5 marker Ex5 (FDR corrected), and NMDA receptor subunit GRIN2C (FDR corrected). The ten cortical maps with the lowest p-values are visualized in Figure 2A.

**Figure 2.**
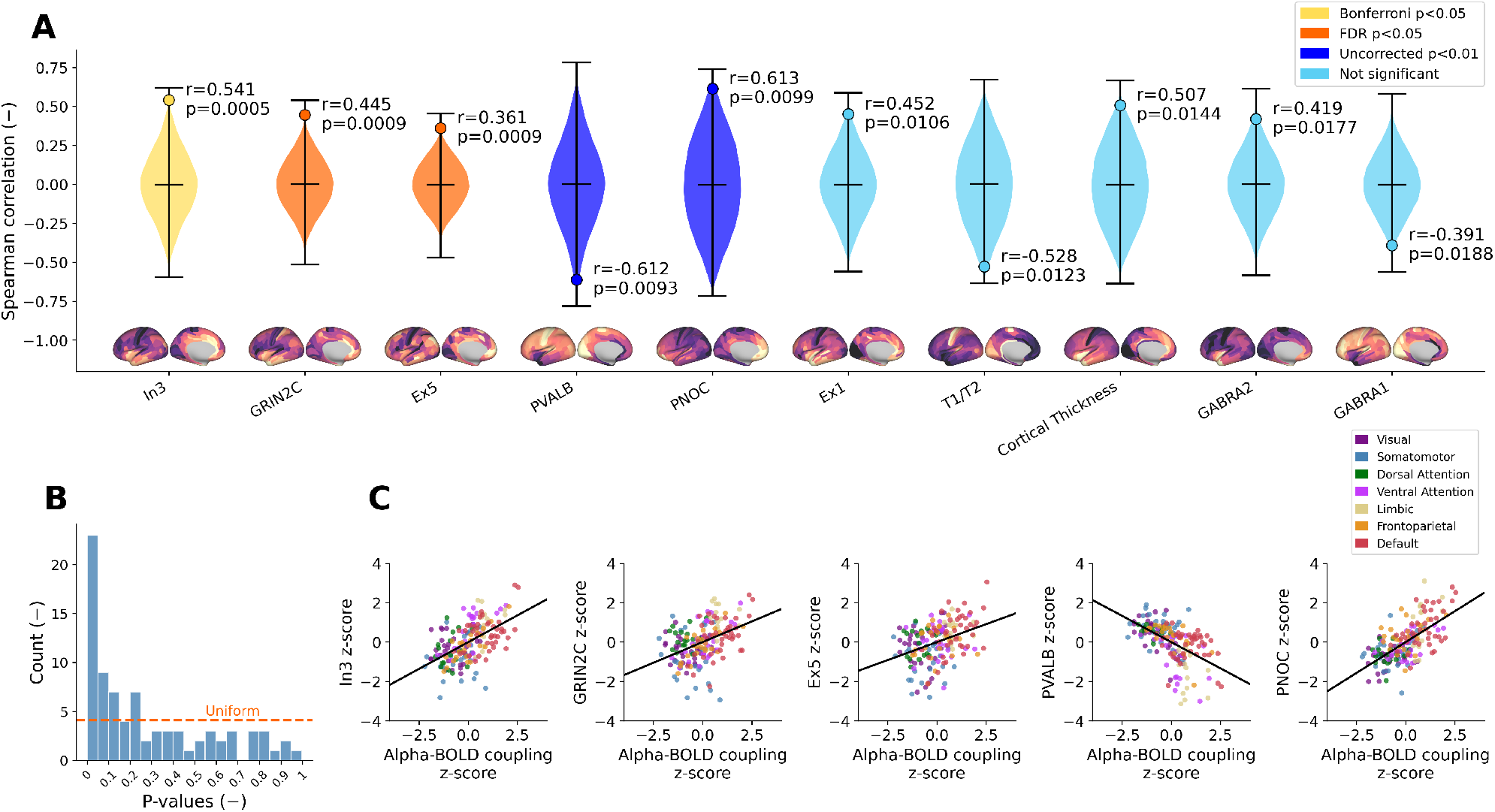
**A**) Violin plots representing the surrogate null distribution for the ten cortical maps with the lowest p-values, together with the observed Spearman correlation (horizontal line). Color codes statistical thresholds: yellow (Bonferroni *p <* 0.05), orange (FDR *p <* 0.05), blue (uncorrected *p <* 0.01), and light blue for the remaining maps. **B**) Histogram of p-values across all 82 tested cortical maps. **C**) Scatter plots with regression lines illustrating the spatial relationships between the alpha-BOLD coupling map and the three FDR-corrected significant cortical maps as well as the two trend-level maps. Points are color-coded by Yeo-7 resting-state network (Yeo et al., 2011).

VIP interneuron marker predominantly points layer 6 VIP interneurons, with a minor fraction also present in layers 1/2 (Lake et al., 2016), and plays a disinhibitory role in cortical circuits (Pi et al., 2013). VIP showed a positive spatial correlation with alpha-BOLD coupling (*ρ* = 0.541, *p* = 0.0005): primary sensory cortices, which have low VIP density, show the most negative alpha-BOLD correlations, whereas associative cortices, with higher VIP density, show less negative or positive coupling.

The excitatory neuron marker reflects layer 5 pyramidal neurons (Lake et al., 2016), which are the major excitatory output cells projecting to other cortical and subcortical regions. This map exhibited a positive correlation (*ρ* = 0.361, *p* = 0.0009): regions with fewer layer 5 pyramidal neurons showed more negative alpha–BOLD correlations.

The GRIN2C map represents NMDA receptor subunit 2C, critical for synaptic plasticity and excitatory transmission. This map showed a positive correlation with alpha-BOLD coupling (*ρ* = 0.445, *p* = 0.0009).

### Two inhibitory interneuron markers show trend-level associations

PVALB showed a negative correlation (*ρ* = − 0.612, *p* = 0.0093): higher parvalbumin interneuron density corresponded to stronger alpha-BOLD anticorrelation. Parvalbumin-positive interneurons are fast-spiking cells known to drive cortical gamma oscillations (Spaak et al., 2012) and to provide powerful perisomatic inhibition of pyramidal cells (Markram et al., 2004).

PNOC (prepronociceptin) marks a less-characterized inhibitory interneuron population (Rodriguez-Romaguera et al., 2020; Ortiz-Juza et al., 2025). Its cortical expression map showed a positive correlation with alpha-BOLD coupling (*ρ* = 0.613, *p* = 0.0099), where regions with lower PNOC interneuron density exhibited more strongly negative alpha-BOLD correlations. The spatial relationships for all five maps are visualized as scatter plots color-coded by resting-state network in Figure 2C.

Notably, several additional cortical maps showed high spatial similarity with the alpha-BOLD coupling map, as reflected in the enrichment of small p-values in Figure 2B. Among these were T1/T2 and cortical thickness. Specifically, T1/T2 serves as an index of myelination (Glasser and Van Essen, 2011) and a reliable proxy for the hierarchical organization of the cortex (Burt et al., 2018), speculatively suggesting that hierarchical organization may be important for reproducing alpha-BOLD coupling in whole-cortex computational models, though this remains to be tested.

### A gene expression model explains alpha-BOLD coupling beyond spatial smoothness

To quantify how well the significant gene maps jointly explain alpha-BOLD coupling, we fitted a multiple linear regression model to rank-transformed data (Figure 3), using the three statistically significant maps (In3, GRIN2C, Ex5) as predictors. The gene-based regression model explained 31.2% of variance in alpha-BOLD coupling across the 180 MMP parcels (*R*^2^ = 0.312, *p* = 0.0003, surrogate median *R*^2^ = 0.052), confirming a highly significant spatial relationship beyond what is expected from spatially autocorrelated noise. For comparison, an analogous structure-based regression model using T1/T2, cortical thickness, and V1 geodesic distance as predictors explained nominally more variance (*R*^2^ = 0.353) but did not reach significance (*p* = 0.187), because the spatial autocorrelation structure of MRI-derived maps closely matches that of the alpha-BOLD coupling map. The variogram panels in Figures 3C and 4C illustrate this directly: structural maps (T1/T2, cortical thickness) closely follow the alpha-BOLD coupling variogram, while gene maps such as Ex5 and GRIN2C show elevated semivariance at 20-50 mm of spatial separation, indicating a spatial autocorrelation structure distinct from the alpha-BOLD coupling map, so the surrogate null distributions yield systematically lower correlations with these maps, making the observed fit unlikely to arise by chance.

**Figure 3.**
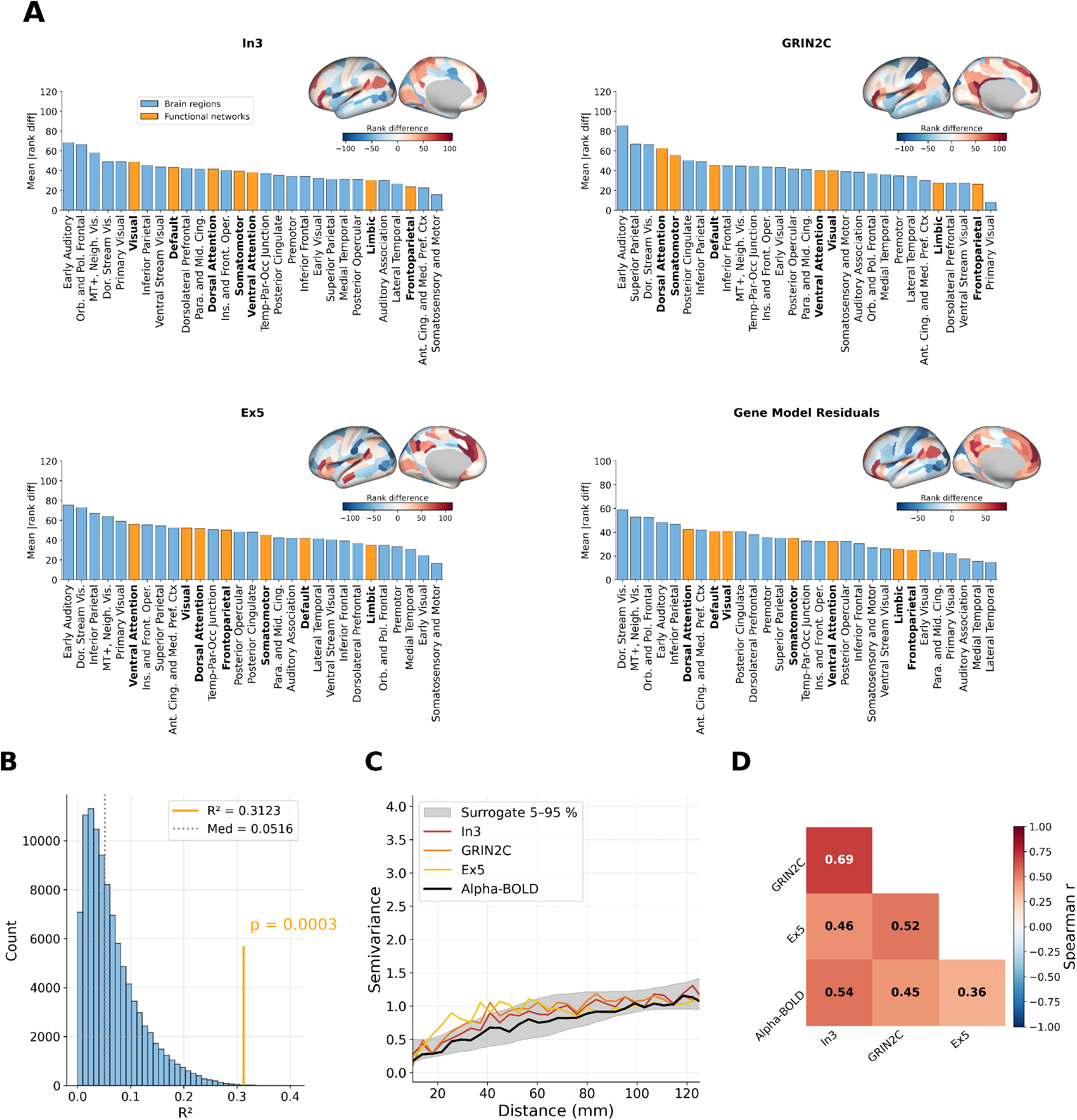
Signed rank differences between the alpha-BOLD coupling map and four gene expression maps. **A**) Each of the four panels shows a brain surface map of signed rank differences (inset, top right; RdBu color scale symmetric around zero: red indicates parcels where alpha-BOLD coupling is more positive than predicted, blue where it is more negative) and a bar chart of mean absolute rank difference grouped by anatomical cortex region (blue bars) and Yeo-7 functional network (bold orange bars), sorted by descending mean. Maps shown: In3, GRIN2C, Ex5, and gene-based regression model residuals. **B**) Gene-based regression model surrogate *R*^2^ null distribution (grey histogram) with the observed *R*^2^ (orange vertical line) and surrogate median (dashed line). **C**) Empirical semivariograms of the three gene expression maps and the alpha-BOLD coupling map (grey shading: 5–95 % surrogate band). **D**) Pairwise Spearman correlation matrix among the three gene expression maps and the alpha-BOLD coupling map.

### Early auditory cortex shows the largest spatial mismatch

In both models (Figure 3), positive values (red on the brain maps) indicate parcels where alpha-BOLD coupling is more positive than predicted by the cortical map, and negative values (blue) indicate parcels where it is more negative than predicted. Early auditory cortex showed the largest rank differences across all three individual gene maps as well as the T1/T2 map, making it the single region where neither molecular nor myelination-based gradients align with the observed alpha-BOLD coupling map.

### Alpha-BOLD coupling is not a simple distance-decay from occipital cortex

The large rank differences for V1 geodesic distance in premotor, inferior frontal, and posterior cingulate cortex suggest that parieto-occipital alpha coupling to BOLD is not a localized occipital phenomenon that gradually fades with cortical distance from V1. If alpha-BOLD coupling were merely an occipital effect attenuating with distance, V1 geodesic distance would be among the strongest structural predictors. Its pronounced failures in frontal and posterior cingulate regions instead suggest that alpha oscillations may engage a spatially selective, network-level coupling pattern consistent with underlying cortical organization rather than simple proximity to occipital cortex.

### Gene- and structure-based maps show structured differences in predicting alpha-BOLD

For the gene model, somatosensory and motor cortex showed the smallest rank differences (Figure 3), while at the network level, the Dorsal Attention and Visual networks showed the largest rank differences, and Frontoparietal and Limbic networks were consistently well aligned. The structural model showed a different pattern: posterior cingulate and premotor cortex showed the largest rank differences, and Somatomotor and Default networks diverged most from model predictions (Figure 4). This dissociation indicates that molecular and structural gradients each fail in distinct large-scale cortical systems, suggesting they might capture different aspects of the spatial organization of alpha-BOLD coupling.

**Figure 4.**
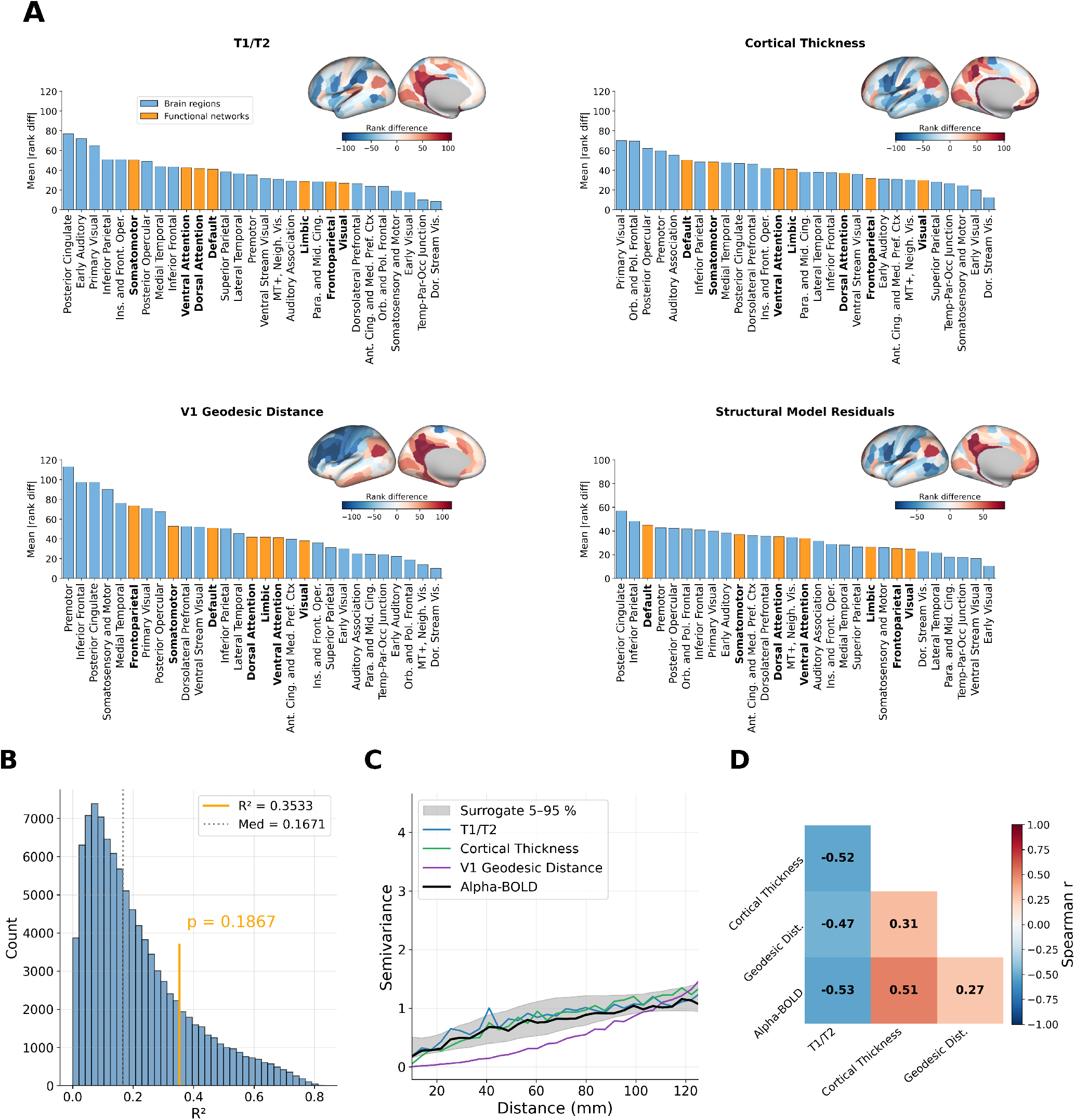
Signed rank differences between the alpha-BOLD coupling map and four structural cortical property maps. **A**) Each of the four panels shows a brain surface map of signed rank differences (inset, top right; RdBu colour scale symmetric around zero: red indicates parcels where alpha-BOLD coupling is more positive than predicted, blue where it is more negative) and a bar chart of mean absolute rank difference grouped by anatomical cortex region (blue bars) and Yeo-7 functional network (bold orange bars), sorted by descending mean. Maps shown: T1/T2 ratio, cortical thickness, V1 geodesic distance, and structure-based regression model residuals. **B**) Structure-based regression model surrogate *R*^2^ null distribution (grey histogram) with the observed *R*^2^ (orange vertical line) and surrogate median (dashed line). **C**) Empirical semivariograms of the three structural cortical property maps and the alpha-BOLD coupling map (grey shading: 5–95 % surrogate band). **D**) Pairwise Spearman correlation matrix among the three structural maps and the alpha-BOLD coupling map.

## Discussion

We compared the resting-state alpha-BOLD coupling map (Jiricek et al., 2025) to cortical maps of gene expression markers for different cellular and receptor properties, as well as MRI structural measures Burt et al. (2018). Three maps showed statistically significant associations: VIP interneuron marker (In3, Bonferroni corrected) (Pi et al., 2013), excitatory layer 5 marker Ex5, and NMDA receptor subunit GRIN2C (Ravikrishnan et al., 2018) (*q <* 0.05, FDR corrected). Two additional inhibitory interneuron markers, PVALB (Ruden et al., 2021) and PNOC (Harris et al., 2018), showed trend-level associations (*p <* 0.01, uncorrected).

The significant correlation with VIP (In3), together with trend-level associations with PVALB and PNOC, highlights the potential role of inhibitory circuits in alpha-BOLD coupling. VIP interneurons (predominantly in layer 6, with a smaller fraction in layers 1/2) operate through disinhibition: by inhibiting SST and other interneurons, they reduce tonic inhibitory tone on pyramidal cells (Pi et al., 2013). Areas with sparser VIP populations may therefore sustain stronger net inhibition during the alpha state, potentially deepening the alpha-BOLD anticorrelation. VIP density thus represents a testable parameter in computational models of alpha-BOLD coupling (Moreni et al., 2025).

Parvalbumin-positive (PV) interneurons, known for their fast-spiking properties, are critical for generating gamma oscillations (Sohal et al., 2009) that positively correlate with BOLD signals in anesthetized monkeys (Logothetis et al., 2001). However, gamma oscillations are less visible in scalp EEG, limiting reliable macroscopic detection under resting-state condition. Gamma and alpha oscillations show a complex relationship, with gamma amplitude phase-locked to alpha and the two rhythms anticorrelating in power (Spaak et al., 2012), suggesting that alpha-BOLD anticorrelation may reflect alpha’s inhibitory role in suppressing gamma-driven excitatory processes in PV circuits. Complementing this, PV interneurons are metabolically demanding, owing to their high tonic firing rates and large number of perisomatic synapses (Markram et al., 2004). We hypothesize that when alpha oscillations suppress PV-driven activity, the reduction in this energetically expensive cycle might produce a disproportionately large BOLD signal drop, offering a metabolic account complementary to the gamma-alpha anticorrelation phenomenon. In frontal areas, where PV density is lower, these effects may be less prominent. This hypothesis is speculative and rests solely on the spatial co-occurrence reported here; direct testing would require simultaneous recordings of PV interneuron activity and metabolic BOLD responses.

The trend-level association with PNOC points to a speculative but interpretable pattern. In rodents, PNOC-expressing neurons in the bed nucleus of the stria terminalis encode rapid arousal changes and responses to salient stimuli (Rodriguez-Romaguera et al., 2020; Ortiz-Juza et al., 2025). If cortical PNOC interneurons serve an analogous arousal-gating role, associative regions with higher PNOC density may be more susceptible to arousal-driven fluctuations that transiently over-ride the resting alpha state, making BOLD dynamics less stably entrained to the alpha rhythm. Sensory regions with lower PNOC density may remain more tightly governed by global alpha-inhibitory rhythms, producing a stronger anticorrelation. This parallels the VIP disinhibition account and should be treated as a hypothesis for future experimental testing, which could be tested in neuronal-type-rich computational modeling frameworks (Moreni et al., 2025).

The Ex5 cluster marks layer 5 pyramidal neurons, for which contradictory views exist on their role in alpha oscillations: they have been proposed as primary generators (Silva et al., 1991), yet laminar LFP studies report stronger alpha associations with supragranular layers (Halgren et al., 2019). Beyond oscillation generation, layer 5 pyramidal neurons are major contributors to local metabolic activity, so regional differences in Ex5 density may be jointly associated with both alpha power and the associated BOLD response, though the directionality of this relationship is not straightforward and should be tested in detailed computational models or experimentally.

High GRIN2C expression aligns spatially with positive alpha-BOLD coupling, suggesting that NMDA receptor signaling may contribute to alpha-related neurovascular dynamics (Attwell et al., 2010). Regional differences in GRIN2C density could be associated with the spatial variation in coupling strength, with regions of lower GRIN2C expression corresponding to more negative alpha-BOLD correlations.

The gene-based regression model reached significance (*R*^2^ = 0.312, *p* = 0.0003) against spatial autocorrelation surrogates. The structure-based regression model explained nominally more variance (*R*^2^ = 0.353) but did not reach significance (*p* = 0.187). This asymmetry is explained by the variogram structure visible in Figures 3C and 4C. The variogram quantifies how semi-variance grows with spatial separation between parcels. The T1/T2 and cortical thickness variograms closely track the alpha-BOLD coupling variogram, meaning BrainSMASH surrogates (constructed to reproduce the same spatial autocorrelation structure) also correlate well with structural maps by construction, making the apparent *R*^2^ reproducible by chance. By contrast, Ex5 and GRIN2C show markedly higher semivariance at 20–50 mm distances, reflecting a spatial autocorrelation structure distinct from alpha-BOLD coupling. Surrogates therefore correlate less strongly with those gene maps, so the observed fit is unlikely to arise by chance.

Compared to MRI-derived metrics such as T1/T2 ratio, cortical thickness (Mahjoory et al., 2020; Burt et al., 2018; Glasser and Van Essen, 2011), gene expression maps offer greater biological specificity, resolving individual cell types, receptor subtypes, and laminar populations rather than broad gradients. This specificity allows identification of concrete biological parameters for incorporation into computational models (Deco et al., 2021; Kong et al., 2021; Sip et al., 2023).

Beyond overall correspondence, we examined the regions where the model’s predictions are weakest. Spatial rank differences pinpoint areas where molecular gradients or structural maps have reduced predictive power for the observed alpha-BOLD organization, suggesting that local network dynamics, state-dependent modulation, or long-range connectivity dominate over fixed anatomical properties. The strongest mismatch across gene maps and T1/T2 involves the early auditory cortex, which all those maps predict should show strongly negative alpha-BOLD coupling alongside other primary sensory regions. The measured coupling, however, is not strongly negative. Two factors may contribute: the cortical surface area of early auditory cortex is roughly ten times smaller than other primary sensory cortices such as visual and somatosensory cortex, or even might be temporarily out of phase, potentially reducing its weight in the group-level parieto-occipital alpha pattern, and the continuous MRI scanner noise likely maintains auditory cortex in a sustained processing state that disrupts resting alpha fluctuations (López-Madrona et al., 2024), decoupling it from the sensory-to-association gradient that molecular and structural maps capture. Cortical thickness maps place the early auditory cortex somewhat higher in the hierarchy, which is why it does not rank as an extreme outlier under that particular map.

At the network level, gene model prediction is weakest in the Dorsal Attention and Visual networks, where alpha-BOLD coupling probably reflects fluctuating attentional state rather than being strongly shaped by cellular composition (Wang et al., 2016; Proskovec et al., 2018). Frontoparietal and Limbic networks are consistently well predicted. The structural model shows a different pattern: prediction is weakest in the Default Mode network, more specifically the Posterior Cingulate Cortex (PCC). Alpha oscillations are linked to positive BOLD in the DMN and negative BOLD in sensory and task-positive networks (Mantini et al., 2007), reflecting a whole-brain anticorrelation dynamic (Fox et al., 2005) in which the two network systems fluctuate in opposition. The alpha-BOLD coupling map might therefore be partly a readout of this network-level anticorrelation: regions with positive coupling (DMN, PCC) are those whose BOLD naturally rises when sensory-network BOLD falls, and vice versa. If so, positive alpha-BOLD coupling at PCC reflects large-scale network dynamics rather than local tissue properties. This would explain why structural features, such as myelination or cortical thickness, which encode regional anatomy rather than network dynamics, systematically underpredict it.

These findings can inform multiple modeling frameworks. Whole-brain network models (Deco et al., 2021; Kong et al., 2021; Sip et al., 2023) could test how spatial variations in cell-type densities and receptor distributions shape large-scale alpha-BOLD coupling. Complementarily, single-area models (Chien et al., 2025; Neymotin et al., 2020; Hahn et al., 2022; Jones et al., 2009) could then isolate local contributions by directly manipulating interneuron density or receptor levels.

Beyond the data analysis frameworks used in this study, other approaches have examined alpha-BOLD coupling at finer spatial resolutions. One notable example is laminar EEG-fMRI (Scheeringa et al., 2016; Scheeringa and Fries, 2019; Clausner et al., 2025), which distinguishes activity across cortical layers. Those studies specifically designed their acquisition protocols to achieve sub-millimeter spatial resolution (0.75 mm isotropic voxels), and used interpolation methods to obtain depth-resolved BOLD estimates. The data used in this study were acquired with a standard whole-brain multiband EPI sequence (3 mm slice thickness), a resolution chosen to maximize temporal resolution and whole-cortex coverage, but insufficient for laminar separation. Nevertheless, laminar EEG-fMRI offers improved alignment with the cortex’s layered structure, providing valuable layer-specific insights into EEG and BOLD signal dynamics.

Another promising avenue involves examining the role of subcortical structures, particularly the thalamus, in the alpha-BOLD relationship as suggested by some (Pang and Robinson, 2018). Although the thalamus is not directly represented in the gene maps analyzed here, certain cortical maps may indicate stronger thalamocortical coupling in specific regions (e.g., areas enriched in layer 6 VIP interneurons).

It is worth noting some uncertainty concerning the group spatial maps themselves. Alpha-BOLD coupling studies using EEG-fMRI (Goldman et al., 2002; Moosmann et al., 2003; Laufs et al., 2003a,b; Tyvaert et al., 2008; de Munck et al., 2009; Gonçalves et al., 2006; Feige et al., 2005) consistently report a broadly similar spatial pattern following the sensory-to-association cortical gradient, yet show slight variations in details, likely due to methodological differences in preprocessing, frequency definitions, and statistics. These differences complicate precise cross-study comparisons. In our study, we employed data-driven spatio-spectral decomposition (Jiricek et al., 2025), a method that is less constrained by the prior selection of an electrode subset or a frequency band of interest than typical approaches, producing reliable alpha oscillation patterns that emerge naturally from the data.

Beyond methodological variability, other sources exist. For instance, intersubject variability in EEG-fMRI is substantial (Gonçalves et al., 2006), as is inter-dataset variability and natural biological variability across individuals and studies. Furthermore, even within the same scanning session, the alpha-BOLD coupling may vary over time, meaning that the session-averaged pattern may represent a mixture of distinct instantaneous states (Mayhew and Bagshaw, 2017). One potential driver of such non-stationarity is vigilance, which acts as a slower, gradual component and can shape the spatial pattern of alpha-BOLD coupling (Olbrich et al., 2009). While averaging smooths out individual differences that may carry important insights, it provides a robust estimate of long-term spatial patterns, which are likely encoded in cortical structure and thus more relevant to cortical maps. The lack of individual cortical maps limits our ability to assess how intersubject variability in spatial features relates to alpha-BOLD coupling variability. Specifically, the transcriptomic cortical maps derived from the Allen Human Brain Atlas (Hawrylycz et al., 2012) are approximate, based on only six post-mortem donors, with tissue samples predominantly drawn from the left hemisphere and collected in a spatially non-uniform manner.

The statistical model used in this study, based on the BrainSmash tool approach (Burt et al., 2020), addresses spatial autocorrelation in brain maps but does not fully capture local inhomogeneities. While it controls for global spatial dependencies, finer-scale patterns may be underrepresented in the null distributions, potentially inflating false-positive rates. Although many cortical maps are spatially similar, our multiple linear regression model represents a first step toward a multivariate account of alpha-BOLD coupling. More advanced multivariate approaches such as regularised regression or dimensionality reduction methods, could help identify combinations of features that better explain the spatial variance, at the cost of added complexity and reduced interpretability of individual feature contributions. Still, pursuing such methods may offer a valuable path for future work, allowing deeper insight into the relationship between cortical architecture and EEG-BOLD coupling.

To conclude, by identifying candidate inhibitory and excitatory neuronal populations and receptor subunits whose spatial expression aligns with alpha-BOLD coupling, this study provides a biologically grounded starting point for future experimental and computational work aimed at understanding how cortical circuits shape this phenomenon.

## Supplementary Note 1: Acknowledgements

The publication was supported by the Czech Science Foundation project No. 21-32608S, the ERDF-Project Brain dynamics, No. CZ.02.01.01/00/22_008/0004643, a Lumina-Quaeruntur fellowship (LQ100302301 awarded to H.S.) by the Czech Academy of Sciences, and the Czech Technical University Internal Grant Agency - grant number SGS23/120/OHK3/2T/13. We acknowledge the core facility MAFIL supported by the Czech-BioImaging large RI project (LM20181291026 funded by MEYS CR) for their support with obtaining scientific data presented in this paper. The cortical maps were derived based on datasets provided by the Human Connectome Project, WU-Minn Consortium (Principal Investigators: David Van Essen and Kamil Ugurbil; 1U54MH091657) funded by the 16 NIH Institutes and Centers that support the NIH Blueprint for Neuroscience Research; and by the McDonnell Center for Systems Neuroscience at Washington University.

## Supplementary Note 2: Authors contributions

**Stanislav Jiricek:** Data curation, Formal analysis, Investigation, Methodology, Software, Validation, Visualization, Writing – original draft, Writing – review & editing. **Vincent S.C. Chien:** Data curation, Formal analysis, Investigation, Methodology, Software, Validation, Visualization, Writing – original draft, Writing – review & editing. **Helmut Schmidt:** Conceptualization, Methodology, Resources, Supervision, Writing – original draft, Writing – review & editing. **Vlastimil Koudelka:** Conceptualization, Formal analysis, Investigation, Methodology, Resources, Supervision, Validation, Writing – original draft, Writing – review & editing. **Radek Marecek:** Data curation, Methodology, Software, Writing – original draft, Writing – review & editing. **Dante Mantini:** Methodology, Software, Writing – original draft, Writing – review & editing. **Jaroslav Hlinka:** Conceptualization, Funding acquisition, Investigation, Methodology, Project administration, Resources, Supervision, Validation, Writing – original draft, Writing – review & editing.

